# *Streptococcus salivarius* - a potential salivary biomarker for orofacial granulomatosis and Crohn’s disease?

**DOI:** 10.1101/422865

**Authors:** Rishi M. Goel, Erica M. Prosdocimi, Ariella Amar, Yasmin Omar, Michael P. Escudier, Jeremy D. Sanderson, William G. Wade, Natalie J. Prescott

## Abstract

**Objective:** Orofacial granulomatosis (OFG) is a rare disease characterised by chronic, non-caseating, granulomatous inflammation primarily affecting the oral cavity. Histologically, it is similar to Crohn’s disease (CD) and a proportion of patients have both OFG and CD. The cause of OFG remains elusive but it has been suggested that microbial interactions may be involved. The aim of this study was to compare the salivary microbial composition of subjects with OFG and/or CD and healthy controls.

**Design:** 261 subjects were recruited, of whom 78 had OFG only, 40 had both OFG and CD, 97 had CD only with no oral symptoms and 46 were healthy controls. Bacterial community profiles were obtained by sequencing the V1-V3 region of the 16S rRNA gene.

**Results:** There were no differences in richness or diversity of the salivary bacterial communities between patient groups and controls. The relative abundance of the *Streptococcus salivarius*-group were raised in patients with OFG or CD only compared to controls while that of the *Streptococcus mitis* -group was lower in CD compared to both OFG and controls. One *S. salivarius* oligotype made the major contribution to the increased proportions seen in patients with OFG and CD.

**Conclusion:** The salivary microbiome of individuals with OFG and CD was similar to that found in health although the proportions of *S. salivarius*, a common oral *Streptococcus* were raised. One specific strain-level oligotype was found to be primarily responsible for the increased levels seen.

## Introduction

Orofacial granulomatosis (OFG) is a rare chronic disease characterised by lip swelling and oral inflammation. The term was originally used to describe oral signs clinically and histologically resembling Crohn’s disease (CD) in patients with no apparent disease elsewhere in the gastrointestinal tract [1]. CD is a chronic, granulomatous, inflammatory condition which can affect any part of the gastrointestinal tract, but most commonly occurs in the terminal ileum. As the mouth is continuous with the gastrointestinal tract and can be affected by granulomatous inflammation, the term “oral CD” has also been used to describe patients with granulomatous inflammation of the oral cavity.

In addition, whilst a majority of patients with OFG without gut symptoms have been shown to have microscopic intestinal granulomata, only a relatively small proportion develop gut CD [2] [3]. Being a rare disorder, the reported geographical distribution of OFG may be skewed by differences in disease classification and reporting. Notwithstanding this, the majority of cases have been reported in the United Kingdom, particularly Scotland, and OFG appears to occur in greater frequency with concurrent CD in Northern Europe as compared to the South [4]. Males and females appear to be affected equally with the median age of disease onset being 23 years [2].

The clinical features of OFG include recurrent lip swelling and oral ulceration. Persistent inflammation can result in disfiguring fibrotic disease which in some cases causes permanent lip swelling refractory to medical therapy which requires debulking surgery. Other features include gingival erythema [5], mucosal tags and ‘cobblestoning’ caused by buccal oedema [6]. Being facially-disfiguring, the disease carries a significant psychological burden for affected individuals [7].

The underlying cause of OFG remains unknown but it is likely to have a multifactorial aetiology. Patients with OFG have a high incidence of atopy compared to the general population [8, 9], and dietary antigens including cinnamon and benzoate compounds may trigger disease exacerbations. Excluding these antigens from the diet has been found to be effective in controlling OFG in up to 25% of individuals and is the first line intervention in management [10, 11]. Immunohistochemistical analysis of oral biopsies from patients with active OFG identified a novel population of subepithelial dendritic B cells which expressed IgE [12], supporting the concept that an antigen may trigger an immediate hypersensitivity reaction.

Other granulomatous diseases such as sarcoidosis or tuberculosis are known to result from infection by specific microorganisms. It has been suggested that *Mycobacterium avium* s.s. *paratuberculosis* (MAP) might play a similar role in CD but has yet to be definitively proven [13, 14]. Raised levels of antibodies against a mycobacterial stress protein have been found in patients with OFG [15], but MAP has not been detected in OFG lesions [16, 17]. The spirochaete *Borrelia burgdorferi* has also been implicated in OFG based on raised antibody levels to the organism and apparent treatment success with penicillin [18, 19] but this finding was not confirmed in a later study [20].

To date, there is no compelling evidence for the role of a specific infective organism in OFG. It is possible, however, that the initiating event for OFG is an inappropriate immune response to a member or members of the normal microbiota giving rise to inflammation. This would change the local environment by altering oral surfaces and thereby changing colonisation patterns and/or providing serum-derived nutrients which enhance the growth of secondary colonisers. The aim of the study was to use 16S rRNA gene community profiling to determine the composition of the salivary microbiome in patients with OFG only, OFG with concurrent CD and compare this to patients with CD without oral involvement and to healthy controls.

## Methods

### Patients and controls

Patients attending a specialist OFG clinic in the department of Oral Medicine at Guy’s & St Thomas’ Hospitals, London, were recruited over a 2-year period. CD patients and healthy controls were recruited via IBD clinics at Guy’s & St Thomas’ Hospitals as previously described [21]. 261 subjects were recruited for the study: 40 had both OFG and CD (OFG+CD), 78 had oral manifestations only (OFG only), 97 were diagnosed with Crohn’s disease without any oral symptoms (CD only) and 46 were healthy controls (HC). The age of the subjects at the time of collection ranged from 16 to 79 years. Each patient provided informed verbal and written consent.

The main inclusion criteria were a confirmed history of active or inactive OFG and/or CD. The diagnosis of OFG was based on clinical features including lip swelling and typical oral ulceration. Where available, histology results were also used to support the diagnosis. The diagnosis of CD was based on conventional clinical, biochemical, endoscopic, histological and radiological criteria.

All patients underwent an oral examination and the sites of involvement and severity of OFG recorded as part of a standardised oral disease activity score (ODAS). Other oral findings were also recorded, particularly dental disease, active carious disease and other oral mucosal changes. Where possible, patients, with their consent, underwent a basic periodontal examination (BPE) to assess for gingival disease. Details of concomitant medical conditions and antibiotic and immunosuppressant drug treatment, as well as tobacco and alcohol usage were also recorded. For those with CD, details of disease phenotype (Montreal classification), behaviour and surgical history was recorded. The study was approved by the Local Research Ethics Committee (Approval No. 12/YH/0172; Yorkshire & The Humber REC).

### Sample collection

Whole saliva was collected by asking patients/volunteers to spit in a universal container until a minimum volume of at least 1ml had been obtained. Saliva samples were immediately placed on ice and then transferred within 3 hours to a freezer for storage at -70°C. All samples were anonymised and coded.

### Bacterial community profiling

DNA was extracted from the saliva samples by means of the Genelute DNA extraction kit (Sigma-Aldrich) and 16S rRNA genes were amplified by PCR with primers 27F (with the YM modification) and 519R [22, 23]. The primers incorporated a unique barcode and Roche 454 adapters. PCR amplicons were purified, sized, quantified and pooled in equimolar proportions. Emulsion PCR and unidirectional sequencing of the libraries was performed using the Lib-L kit and Roche 454 GS-FLX Titanium sequencer.

### Data analysis

Pre-processing and analysis of sequences was carried out using the mothur analysis suite version1.36.1 [24] based on the Schloss SOP (January 2016). Initial de-noising was performed using AmpliconNoise algorithm. Subsequently any sequences less than 440 bases in length and/or had one of the following; >2 mismatches in the primer, >1 mismatch in barcode regions and homopolymers of >8 bases were removed from the dataset. The remaining sequences were trimmed to remove the primers and barcodes and aligned to the SILVA 16S rRNA reference alignment [25]. An additional pre-cluster step was performed in mothur to merge sequences with four or fewer bases differences. The UChime algorithm [26] as implemented by mothur was used to identify sequence chimeras, which were removed from the analysis. Sequences were clustered into Operational Taxonomic Units (OTUs) at a sequence dissimilarity distance of 0.015 using an average neighbour algorithm and then classified using a Naïve Bayesian classifier implemented in mothur with the Human Oral Microbiome Database reference dataset v13.2 [27]. The α-diversity of bacterial communities based on OTUs was analysed using approaches implemented by mothur: richness of the communities was assessed by the number of observed OTUs and the Chao1 richness index; diversity of the communities was estimated using Simpson’s inverse diversity index. Richness and diversity estimates were compared between groups using Kruskal-Wallis test.

For the β diversity analyses the datasets were normalised by sub-sampling to a level of 3076 sequences per sample, which excluded 19 samples. To compare the β diversity of samples based on OTUs, the thetaYC metric (compares community structure by accounting for the relative abundance of taxa) [28] was used to generate distance matrices in mothur. Analysis of Molecular Variance (AMOVA) [29] as implemented by mothur was then performed to determine if any differences between the microbiomes of our experimental groups were statistically supported by differences in the distance matrix. LeFSE [30] was used to identify OTUs differentially abundant between groups.

### Minimum Entropy Decomposition (oligotyping)

Minimum Entropy Decomposition was performed on the same samples used for the alpha and beta diversity analyses, excluding the 19 samples falling below 3076 sequences. Denoising, alignment, chimera removal and taxonomy assignation were performed using the Mothur analysis suite [24] as described above. Sequences identified as belonging to the genus *Streptococcus* were then extracted and formatted using the “mothur2oligo” tool (available from https://github.com/michberr/MicrobeMiseq/tree/master/mothur2oligo).

Minimum Entropy Decomposition analysis [31] was performed using MED pipeline version 2.1 (available from https://meren.github.io/projects/med/). The MED algorithm is similar to the previously described oligotyping algorithm [32] and differentiates taxa on the basis of single-nucleotide differences in the positions of highest entropy. The parameters used were minimum substantive abundance of a MED node (-M)=7 and maximum variation allowed in each node (-V)=4 nt. The total number of *Streptococcus* sequences analysed was 564,342. Of these, 62,706 were removed as outliers due to the minimum substantive abundance parameter (-M, set to 60) and 8,262 were removed as outliers due to the maximum variation at each node parameter (-V, set to 5). Thus, after the refinement, 493,374 were analysed and classified into 370 MED nodes (oligotypes). Sequences representative of each oligotype were identified at species level by comparison to the HOMD database through the BLAST web tool accessible at http://www.homd.org.

## Results

A total of 1,630,578 sequences were obtained after denoising and quality filtering. Figure 1 shows the bacterial genera found in the samples by group. For most individuals, the communities were dominated by the genera *Streptococcus* and *Prevotella*, although some subjects had no or very few streptococci.

**Figure 1.**
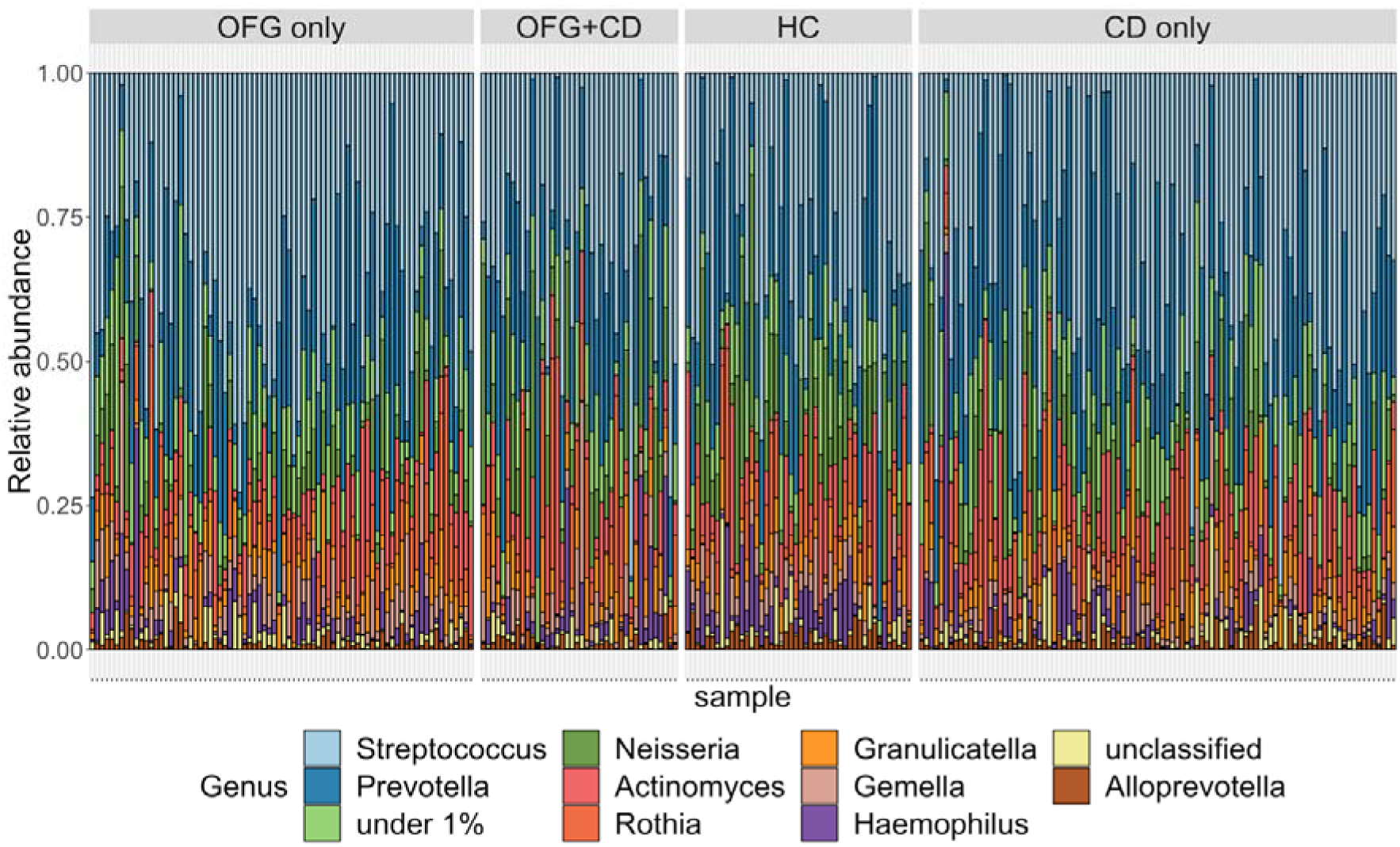
Relative abundance of bacterial genera by patient group. The group “under_1%” combines all genera present at less than 1 % relative abundance.

The alpha-diversity within each sample group was calculated using the Chao1 estimate of richness and the inverse Simpson’s diversity index after subsampling to 3,076 sequences per sample (Table 1). Subsampling at this level removed 19 samples. Subject group sizes for the subsequent analyses were as follows: OFG only (74), OFG+CD (38), CD only (85) and HC (45). There was no significant differences between the groups for the Chao1 or inverse Simpson’s indices, nor for the average number of observed OTUs (Kruskal Wallis test).

**Table 1.** Richness and diversity of the salivary microbiota in subject groups.

| Subject group | n | observed OTUs<br>(sd) | Chao1<br>(sd) | Inverse Simpson<br>(sd) |
| --- | --- | --- | --- | --- |
| OFG only | 74 | 198.6<br>54.2 | 505.1<br>152.5 | 11.2<br>6.2 |
| OFG+CD | 38 | 183.4<br>57.2 | 449.5<br>125.6 | 9.7<br>4.8 |
| HC | 45 | 196.1<br>53.5 | 496.3<br>161.1 | 11.0<br>4.6 |
| CD only | 85 | 203.5<br>57.5 | 493.4<br>133.6 | 9.2<br>5.1 |

The composition of the microbial communities among the four subject groups was significantly different (AMOVA, p-value<0.001). Pairwise comparisons of the individual subject groups were performed and CD appeared to be the primary driver of inter-group differences. For example, the CD only group was significantly different to both the HC group (p<0.001; significance threshold using Bonferroni correction: 0.008) and the OFG only group (p<0.001). The OFG+CD group was significantly different to HC (p=0.006) while OFG only was not significantly different to HC.

As oral disease status is known to be a major factor influencing oral microbiome composition, the basic periodontal examination (BPE) was used to assess the subjects’ gingival health. Total BPE scores were measured for 207 of the 261 subjects and were significantly different between phenotype groups (p<0.001, Kruskal Wallis), as shown in Figure 2. Overall, there was a clear trend for subjects with OFG only to have higher BPE scores than HC whereas CD only patients had lower BPE scores. In view of this, the effect of BPE score on microbiome composition was investigated. BPE scores were assigned to three class variables: low, <= 2; middle, 2-10; high, >= 10. There was no significant difference in microbial composition between BPE class groups by AMOVA.

**Figure 2.**
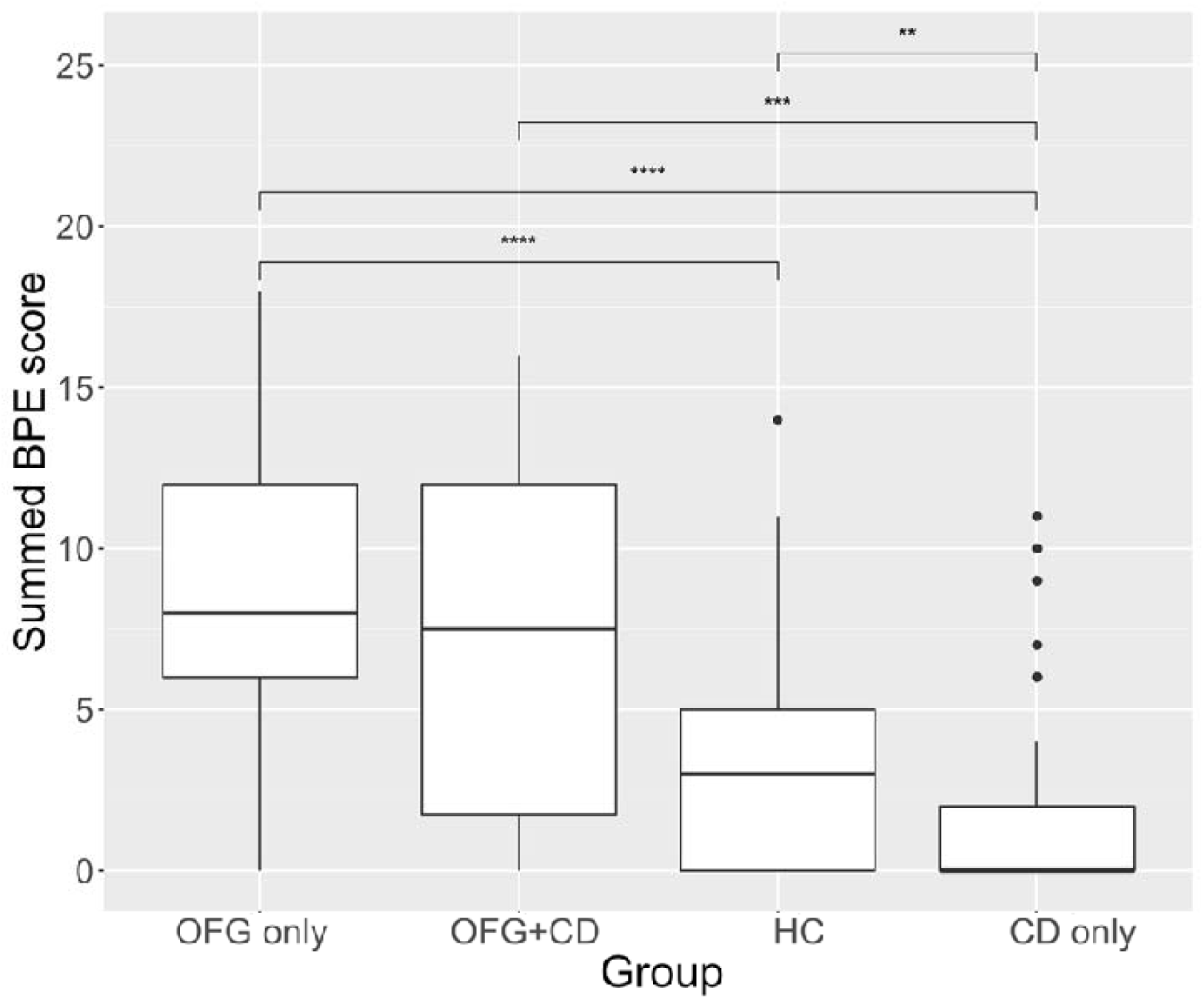
Box plot showing summed BPE scores as a proportion of the total microbiota. Upper and lower edges of the boxes are the first and third quartiles; the line inside the box is the second quartile (median); individual dots are outliers (^∗∗^ - p<0.01; ^∗∗∗^ - p<0.001; ^∗∗∗∗^ - p<0.0001)

LeFSE analysis was used to identify specific OTUs responsible for the differences in microbial composition seen between groups (Table 2). The default threshold of 2 was used for the logarithmic LDA score for discriminative features. Eleven OTUs were found to be differentially represented between groups. Seven of these overrepresented OTUs were overrepresented in the HC group, suggesting that the disease phenotypes were associated with loss of normal microbiota components. Most of the differentially represented OTUs were of relatively low abundance with only three of them (OTU 1, 2 and 6) present in the dataset at a relative abundance of greater than 0.01. OTU 6 was identified as *Haemophilus parainfluenzae* and its relative abundance was significantly reduced in OFG only and CD only compared to HC (Figure 3). OTUs 1 and 2 were identified as unclassified members of the genus *Streptococcus* and were the most frequently detected OTUs in the study, making up 17.3 and 7.8 % respectively of the oral bacterial community across all subjects. Because *Streptococcus* species vary widely in the roles that they play in oral ecology and disease, Minimum Entropy Decomposition (MED) was used to lend more precision to species-level identification.

**Figure 3.**
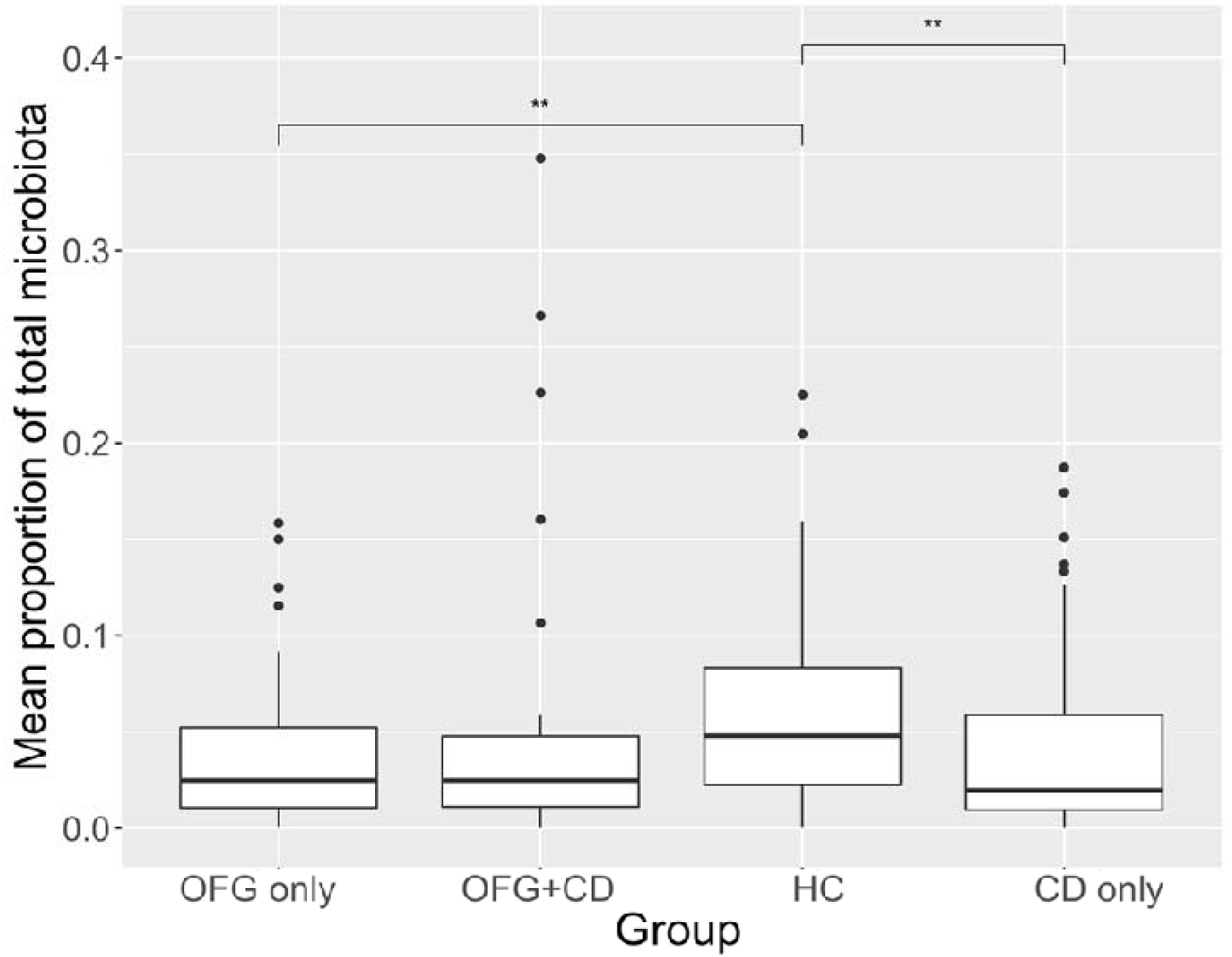
Box plot showing the relative abundance of OTU 6 (*Haemophilus parainfluenzae*) as a proportion of the total microbiota. Upper and lower edges of the boxes are the first and third quartiles; the line inside the box is the second quartile (median); individual dots are outliers (^∗∗^ - P<0.01).

**Table 2.**
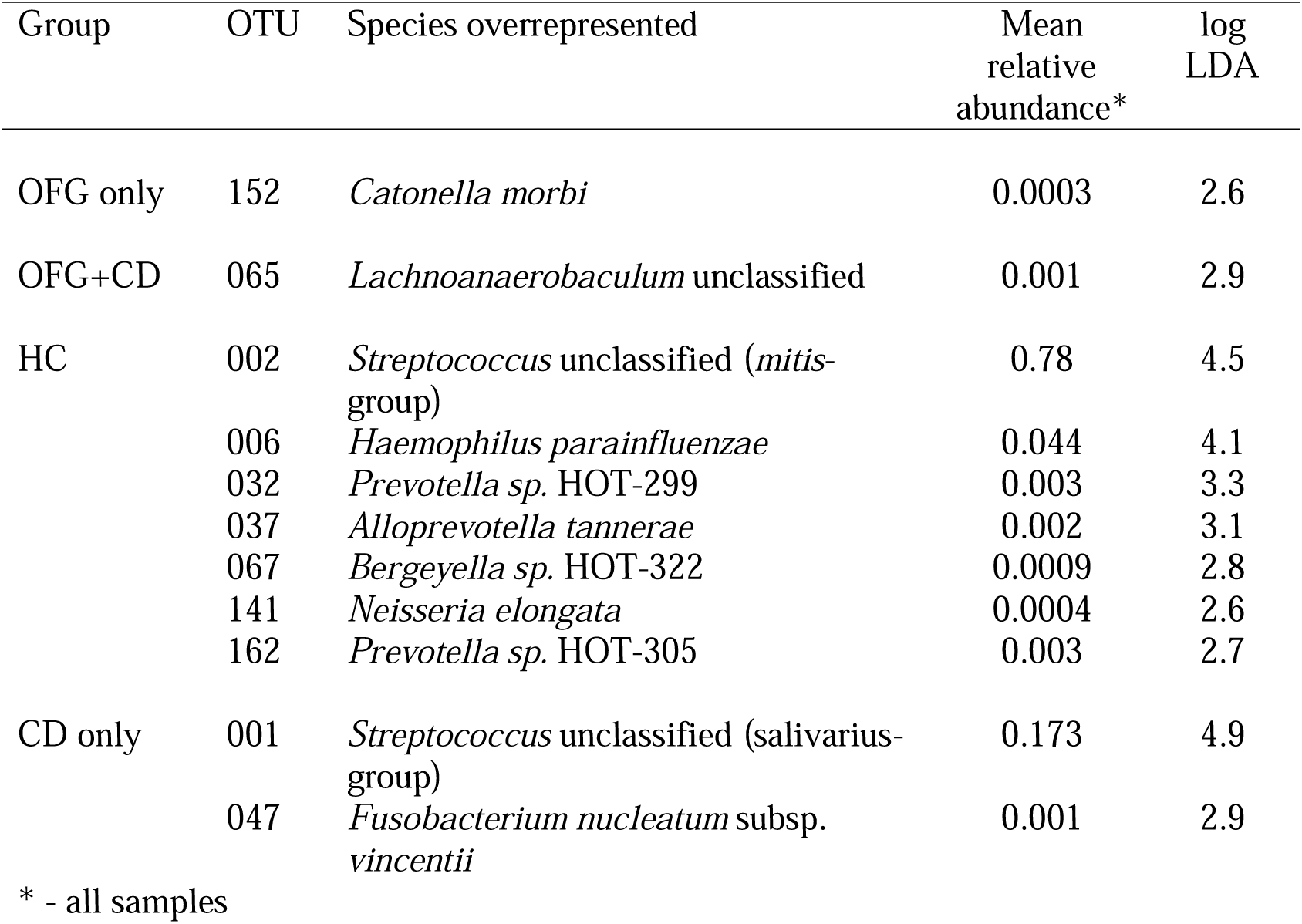
OTUs overrepresented in subject groups (LeFSE).

The *Streptococcus* sequences were binned into 370 oligotypes that were subsequently grouped by species after BLAST interrogation of the HOMD database. Where multiple species-level BLAST identifications were above 98.5% sequence identity, the identification was made to a group of species. The mean abundances of species and species groups making up more than 1% of the total microbiota were compared between groups, using Wilcoxon test with Bonferroni correction for multiple testing. *S. salivarius* proportions were significantly higher in the OFG only and CD only groups compared to HC (Figure 4). The *S. mitis*-group also showed significant differences among groups (Figure 5), with healthy HC and OFG only having significantly higher relative abundances than CD only. In addition, the relative abundances of individual oligotypes whose mean relative abundance was over 0.5% were compared across the groups. Three *S. salivarius* oligotypes were found to show significant differences between groups (Figure 6). Oligotype 2869 showed the largest differences with the OFG only, CD only and OFG+CD groups all having significantly higher relative abundance than the HC group.

**Figure 4.**
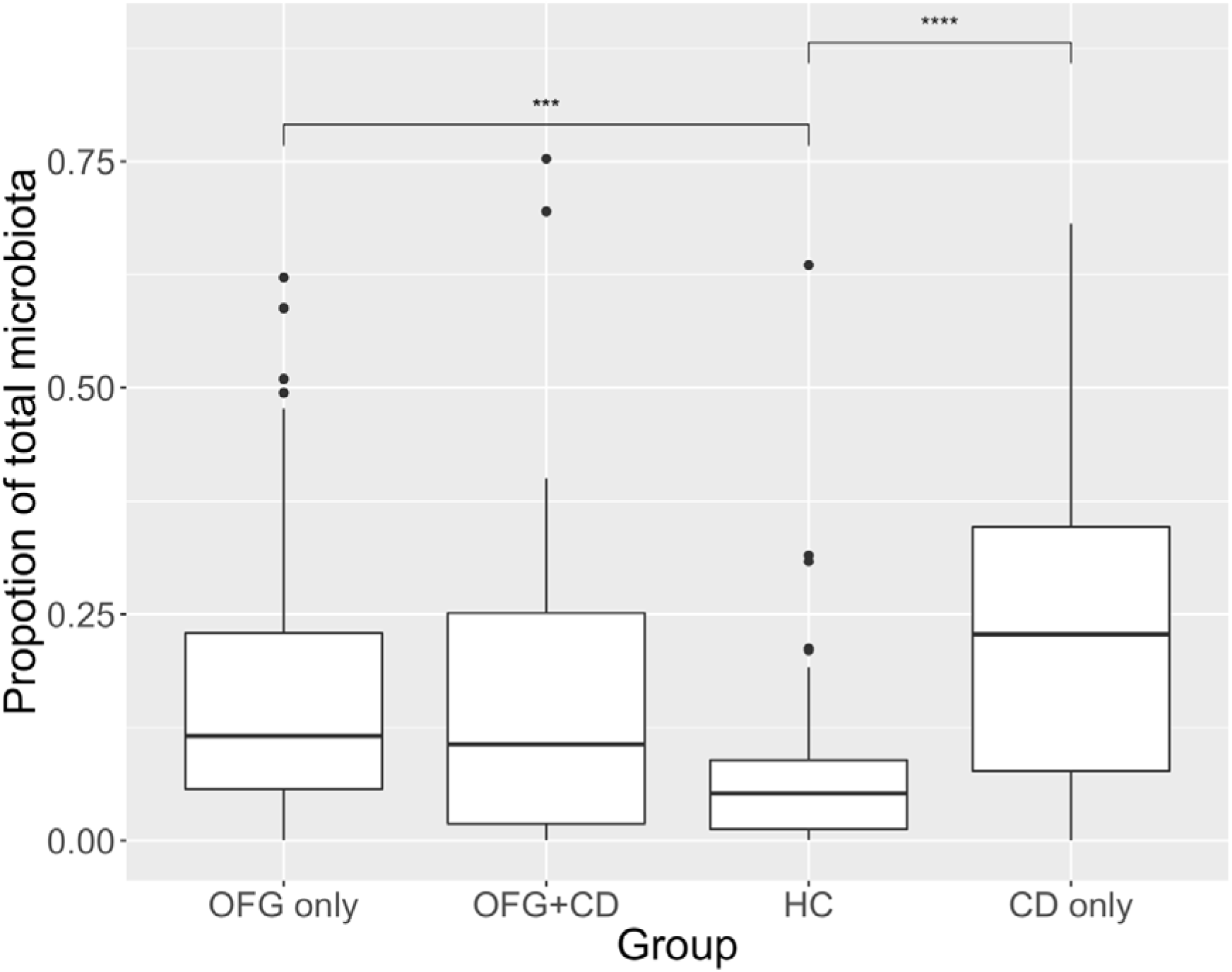
Box plot showing summed *S. salivarius*-group oligotypes as a proportion of the total microbiota. Upper and lower edges of the boxes are the first and third quartiles; the line inside the box is the second quartile (median); individual dots are outliers (^∗∗∗^ - p<0.001; ^∗∗∗∗^ - p<0.0001).

**Figure 5.**
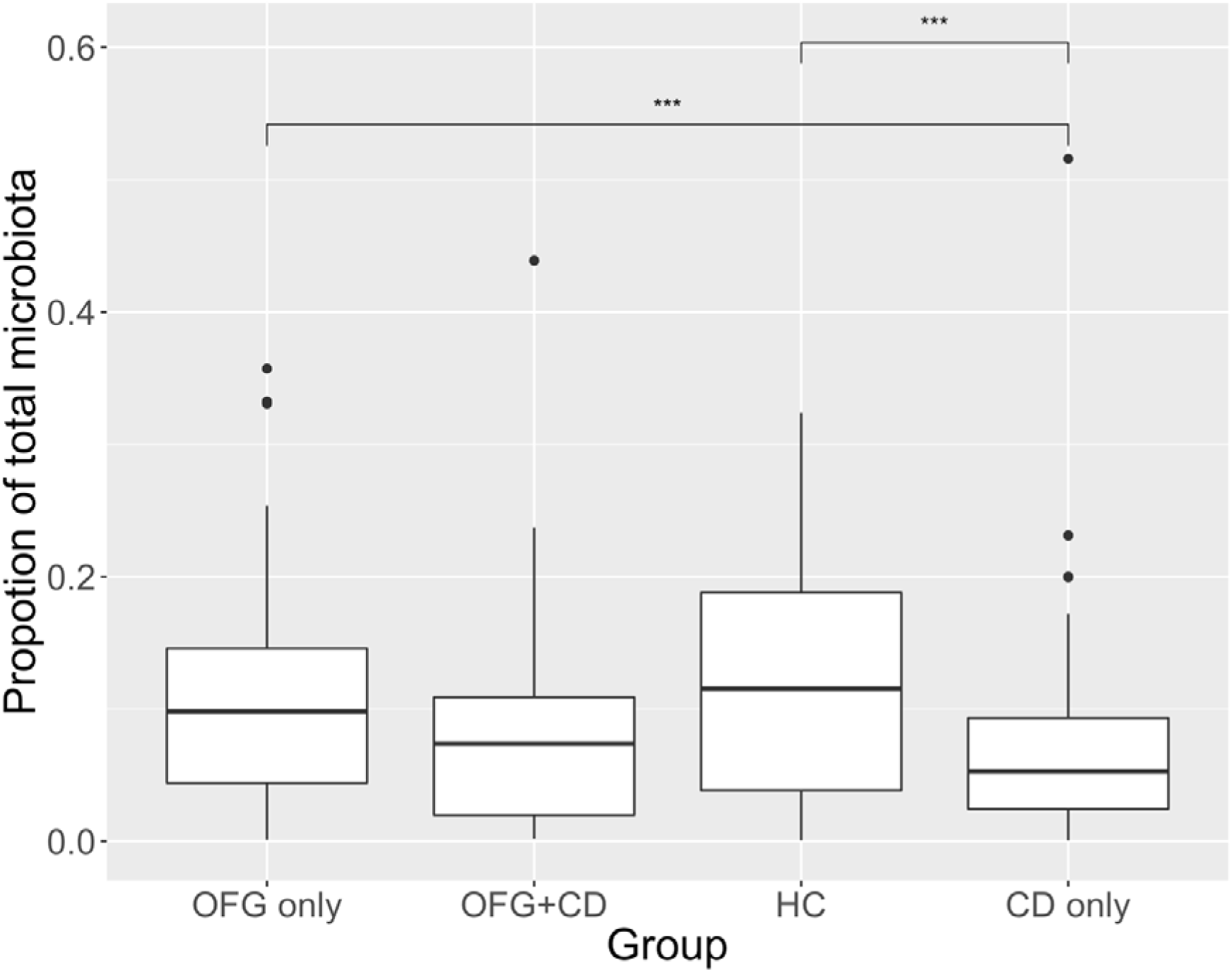
Box plot showing summed *S. mitis*-group oligotypes as a proportion of the total microbiota. Upper and lower edges of the boxes are the first and third quartiles; the line inside the box is the second quartile (median); individual dots are outliers (^∗∗∗^ - p<0.001).

**Figure 6.**
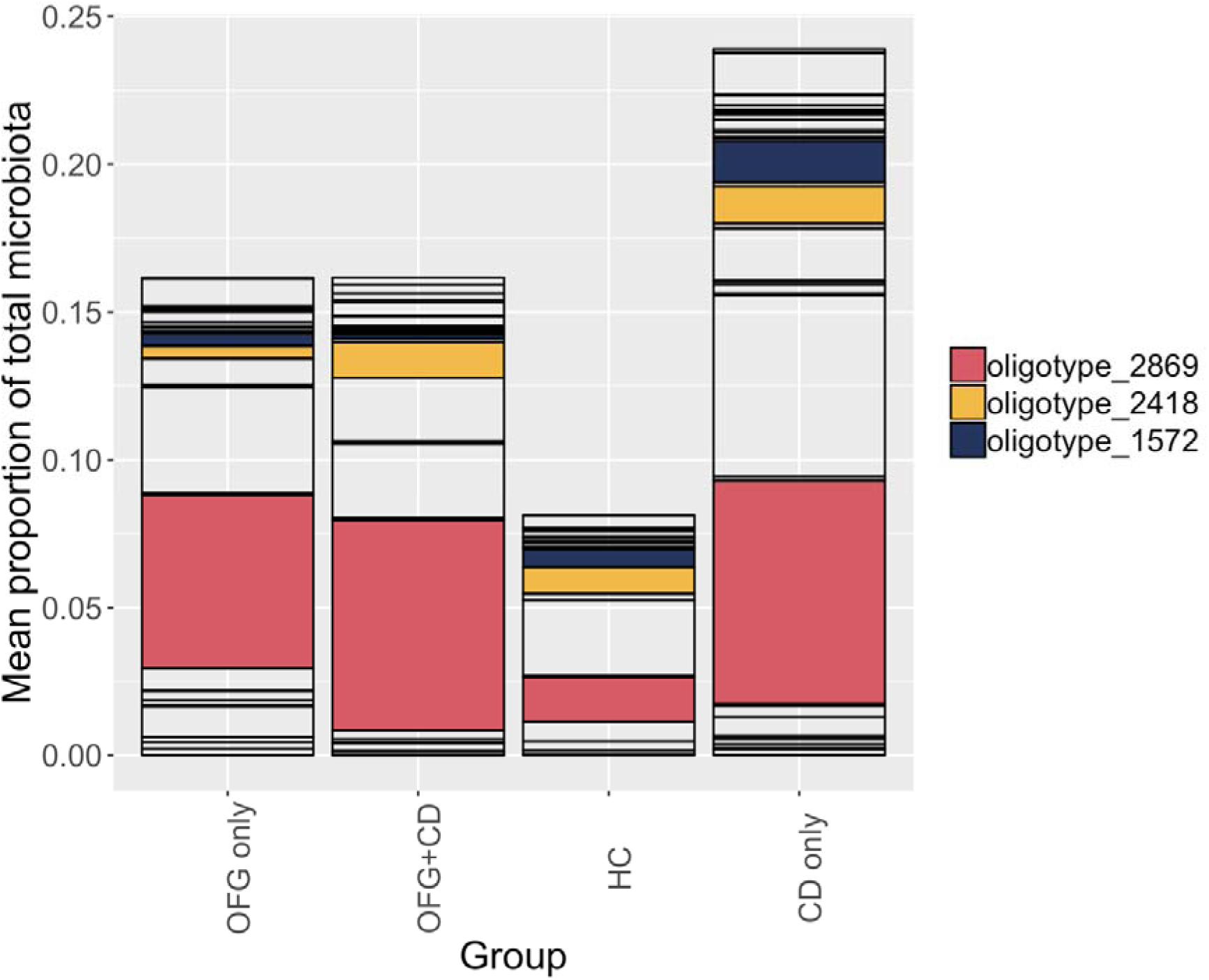
Proportions of individual *S. salivarius*-group oligotypes in subject groups. Oligotypes highlighted in colour showed significant differences between groups (Kruskal-Wallis test).

## Discussion

The results of this study indicate that the oral microbiome in subjects with orofacial granulomatosis is not markedly different to that of healthy controls or subjects with Crohn’s disease because there were no differences in richness or diversity between the groups and all subjects had a typical oral microbiome compositional profile. This finding is in contrast to the results of numerous studies looking at the effect of CD on the composition of the intestinal microbiome [33]. The faecal and mucosal microbiome is substantially altered in patients with CD. Richness is reduced [34, 35] and the phylum *Firmicutes* is relatively depleted particularly anaerobes from the order *Clostridiales*, but raised proportions of *Proteobacteria*, mainly *Enterobacteriaceae*. The shift towards a less anaerobic bacterial community is thought to be the result of increased levels of reactive oxygen species produced as a part of the inflammatory response [34]. Specific genus-level microbial signatures of CD have been reported to be reduced levels of *Faecalibacterium*, an unknown *Peptostreptococcaceae*, *Anaerostipes*, *Methanobrevibacter*, an unknown *Christensenellaceae* and *Collinsella* and increased proportions of *Fusobacterium* and *Escherichia* [36].

Patients with OFG were found to have higher BPE scores than controls although there was no corresponding difference in microbiome composition. The levels of BPE scores seen were indicative of some degree of gingival inflammation and may have been due to poorer oral hygiene in the patients due to the discomfort caused by the OFG lesions.

The lack of substantial alteration of the oral microbiome in OFG with or without gut CD most likely reflects the fact that the oral microbiome is extremely stable and not greatly affected by diet [37] or administration of antibiotics [38]. There were however some OTUs which showed differences in relative abundance between groups in the LEfSe analysis. The high number of comparisons performed in microbiome studies when OTU relative abundances are compared between patient groups can lead to spurious associations being revealed by chance, even when significance thresholds, as here, are corrected for multiple comparisons. Such associations should therefore be interpreted with caution. Of the 11 OTUs over-represented in particular subject groups, 7 were in the control group. This suggests that the major shift in OFG and CD was the relative loss of normal microbiota taxa. Another important consideration in interpreting OTU association analyses is whether the size of the effect is biologically significant. Many of the OTUs found to be differentially abundant were present at extremely low levels and therefore only those present at a relative abundance of greater than 1 % were considered further. Levels of *H. parainfluenzae* were reduced in all patient groups compared to controls. *H. parainfluenzae* is a commonly occurring member of the normal microbiota and the significance of this finding is unclear. In contrast, proportions of oligotypes belonging to the *S. salivarius* group, were found to be significantly raised in the OFG only and CD only groups. *S. salivarius* and related species are regarded as health-associated and are found primarily on the dorsum of the tongue and the pharyngeal mucosa [39, 40]. Indeed strains of *S. salivarius* are used as probiotics with beneficial properties against oral conditions such as halitosis and pharyngitis [41] and have been shown to have anti-inflammatory properties *in vitro*, via down regulation of the NF-κB pathway [42]. It is not clear why the proportions of these species should be raised in OFG and CD but the diseases may change the oral mucosa in ways which promote the adherence and retention of these species. It is particularly interesting that one oligotype, 2869, was specifically elevated in subjects with OFG or CD. It is known that strains of a species can differ markedly in their biological and pathogenic properties and it appears that members of this oligotype, found in multiple subjects, has a particular, and numerically strong, relationship with OFG. These findings are of particular interest as bacterial antigens, including streptococci are a known common target for IgE. And previous studies have identified infiltrates of dendritic B cells in the oral epithelium OFG patients which express surface Immunoglobulin E (IgE) [12]. Future work should be focused on confirming the association of specific *S. salivarius* strains with OFG by metagenomic analyses that enable strain differentiation [43], together with the isolation of representatives of this oligotype and investigation of its properties of relevance to OFG.

In contrast to *S. salivarius*, *S. mitis*-group organisms were present at lower relative abundance in the subjects with CD, compared to OFG and controls. *S. mitis* is the commonest streptococcal species found in the human mouth [40]. The numbers of this group may have appeared to have been reduced because proportions of *S. salivarius* were raised, which will have affected their relative abundance.

The results of this study demonstrate that the overall composition of the salivary microbiota in OFG and CD was similar to that of healthy controls but that there were some significant, and interesting, differences in levels of two of the commonest groups of oral streptococcal commensals which warrant further investigation. In particular, *S. salivarius* was increased in both CD and OFG while *S. mitis* decreased in CD only.

It has been previously shown that pathogenic variants in genes known to confer a high-risk for CD are enriched in OFG patients with concurrent intestinal disease (OFG+CD) [44]. Furthermore, we have recently demonstrated that a useful toolkit for predicting intestinal inflammation in individuals at greatest risk for CD can be implemented from a small set of genetic markers, such as these, combined with family and lifestyle risk factors [45]. The study described here provides the first step in the investigation of the utility of salivary microbial biomarkers as proxy for gastrointestinal dysbiosis and disease. Further studies that can correlate these findings with host genetics, immune status, metabolomics and other risk factors could help to develop prediction tools to identify those OFG patients that are at greatest risk of developing intestinal CD thereby opening up the possibility for early intervention. In addition, a better understanding of the interactions between microbial shifts, inflammation and disease pathology would have the potential to lead to further targets for drug development and disease management strategies for this complex phenotype.

## Acknowledgements

This work was supported by the National Institute for Health Research (NIHR) Biomedical Research Centre at Guy’s and St Thomas’ NHS Foundation Trust and King’s College London and The Wellcome Trust (094491/Z/10/Z). The support of the clinical and nursing staff of the Oral Medicine Unit, Guy’s and St Thomas’ Hospitals is gratefully acknowledged.

